# Flux, toxicity and protein expression costs shape genetic interaction in a metabolic pathway

**DOI:** 10.1101/362327

**Authors:** Harry Kemble, Catherine Eisenhauer, Alejandro Couce, Audrey Chapron, Mélanie Magnan, Gregory Gautier, Hervé Le Nagard, Philippe Nghe, Olivier Tenaillon

## Abstract

Our ability to predict the impact of mutations on traits relevant for disease and evolution remains severely limited by the dependence of their effects on the genetic background and environment. Even when molecular interactions between genes are known, it is unclear how these translate to organism-level interactions between alleles. We therefore characterized the interplay of genetic and environmental dependencies in determining fitness by quantifying ~4,000 fitness interactions between expression variants of two metabolic genes, in different environments. We detect a remarkable variety of environment-dependent interactions, and demonstrate they can be quantitatively explained by a mechanistic model accounting for catabolic flux, metabolite toxicity and expression costs. Complex fitness interactions between mutations can therefore be predicted simply from their simultaneous impact on a few connected molecular phenotypes.

---

Despite its centrality to medical and evolutionary genetics, our ability to predict the impact of mutations on even the apparently simplest of organismal traits (*1–8*), let alone complex ones (*9*), remains minimal. Three of the main factors proposed to account for this “missing heritability” (*9*) are: the large number of possible alleles at any locus, each having a potentially different impact on a gene’s function; interaction between alleles at different loci (intergenic epistasis), such that their combined effect is not simply the sum of their individual effects; and interaction between genotype and environment, such that different genotypes respond to the environment in different ways (*1–9*). A promising inroad is the increasingly refined characterization of molecular interaction networks enabled by –omics approaches (*10*). Metabolic networks are the best-characterized of these, and are of great practical interest for medicine and engineering, but even for metabolic genes it remains unclear how functional interactions at the molecular level translate to allelic interactions at the level of integrated traits relevant for disease, industry and adaptation (*11*).

We therefore developed an experimental system with which to systematically quantify the fitness interactions occurring between many alleles of two metabolic genes from the same pathway. Further, the design enabled us to probe the dependence of these interactions on environmentally modulated gene expression, a common non-genetic mechanism for the modification of physiological traits (*5, 12*).

Our system was composed of the genes (*araA* and *araB*) encoding the enzymes responsible for the first two steps of the well-studied *Escherichia coli* L-arabinose-utilization pathway (*13*): L-arabinose isomerase (AraA) and L-ribulokinase (AraB), who together transform the sugar, L-arabinose, into the intermediate, L-ribulose-5-phosphate (Fig. 1A). L-ribulose-5-phosphate enters the pentose phosphate pathway (PPP) of central metabolism via further enzymatic reactions, ultimately supporting cell growth, but like many intermediates (*14, 15*), its accumulation is toxic, retarding growth (*16*). Environmental modulation of gene expression was achieved by placing each of the two genes under an independent, *trans* regulated chemically-inducible promoter.

**Fig. 1.**
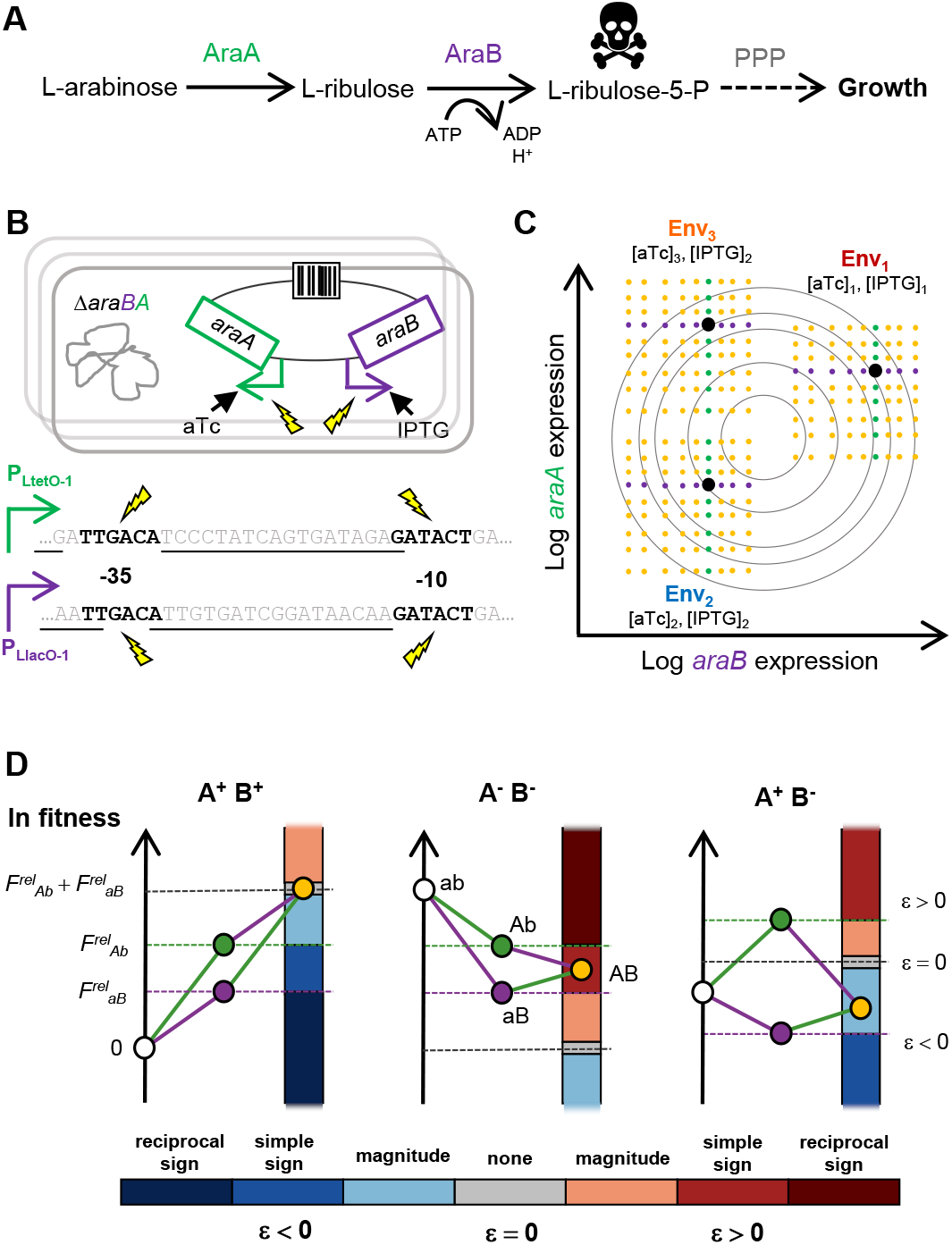
Quantitative mapping of fitness interactions between expression variants of two metabolic genes in expression-modifying environments. **(A)** L-arabinose pathway of *E. coli*. **(B)** *araA* and *araB* were placed under the control of inducible promoters, making their expression sensitive to the concentration of their respective inducers, anhydrotetracycline (aTc) and isopropyl β-D-1-thiogalactopyranoside (IPTG). A barcoded library of mutant promoter combinations was constructed, with mutations targeting the −35 and −10 RNA-polymerase binding hexamers (black letters). Underlined bases are annotated repressor binding sites. **(C)** Competitive fitness was measured under different inducer concentrations defining three environments. P_LtetO-1_ single mutants – green; P_LlacO-1_ single mutants – purple; double mutants – orange. Contours are hypothetical fitness isoclines. **(D)** Epistasis was quantified for all mutant promoter pairs across environments. Epistasis can be categorized as either magnitude or sign type. Sign epistasis is further categorized as simple (effect of one mutation changes sign in presence of the other) or reciprocal (effects of both mutations change sign in the presence of the other). Capitalized letters represent mutant alleles of P_LtetO-1_-*araA* and P_LlacO-1_-*araB*. Superscript plus and minus denote that individual alleles are beneficial or deleterious, respectively.

For each promoter, 36 single-base variants were constructed, along with the initial “wildtype” sequence, and combined with all variants of the other promoter (Fig. 1B). The organismal phenotype, competitive fitness, was then measured for the entire set of 1,369 genotypes under three different inducer concentration combinations (Figs. 1C-D). Fitness was measured by tagging the mutant library with unique DNA barcodes (tens to thousands per genotype) (Figs. S1-2), culturing the pooled library for ~30 mean generations, and tracking barcode frequencies over time with Next-Generation Sequencing (Fig. S3). The barcodes act as internal replicates for every genotype, enabling precise fitness estimates at high-throughput (log relative fitness, *F^rel^*, median standard deviation of 0.0011 for single mutants and 0.0047 for double mutants; Fig. S4).

The overall distribution of fitness effects depended critically on the inducer environment, *ie*. the *trans*-regulatory input (Fig. 2A; Fig. S5A; Data S1). The proportion of beneficial effects varied from 88% in Env_1_ (median *F^rel^* = 0.12) to 51% in Env_3_ (median *F^rel^* = −0.03) and 12% in Env_2_ (median *F^rel^* = −0.12). Further, the correlation of fitness effects between environments ranged from strongly positive (Env_1_-Env_3_, Pearson’s r = 0.74, p < 2.2×10^−16^) to weakly negative (Env_1_-Env_2_, Pearson’s r = −0.11, p = 1×10^−4^) (Fig. S5B), demonstrating that fitness in one environment can be an extremely poor predictor of fitness in other environments due simply to expression differences. At the level of individual alleles, all but one had changing patterns of effects across environments (Fig. 2B). In some environments, they were universally beneficial or deleterious across genetic backgrounds, and in others they switched between being beneficial and deleterious depending on the allele at the second promoter. This pervasive and inconsistent variability poses a clear challenge for the prediction of mutation effects.

**Fig. 2.**
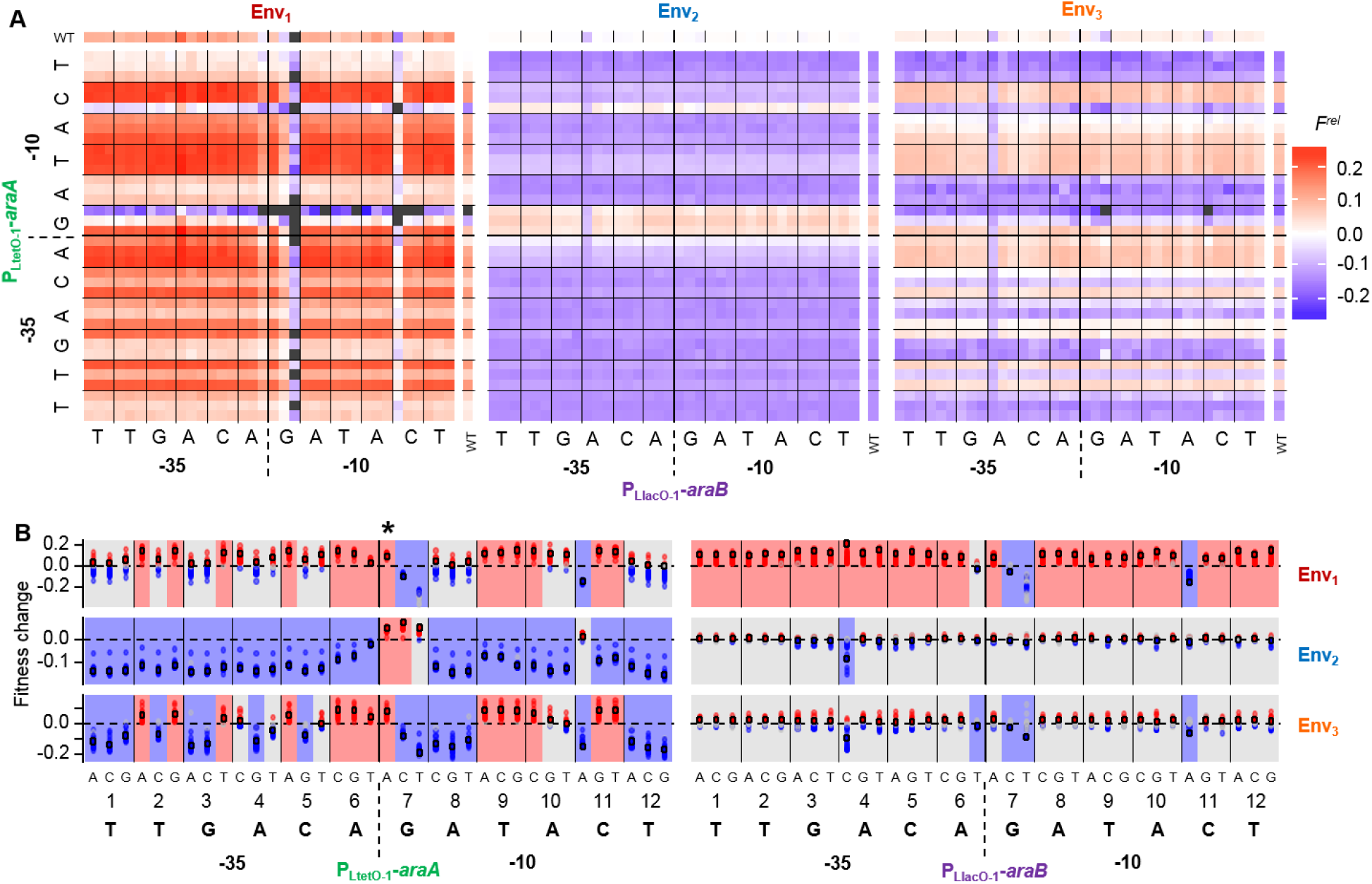
Fitness effects of promoter mutations across backgrounds and environments. **(A)** Genotypes are colored according to the natural logarithm of their fitness relative to the wildtype (*F^rel^*). Grey denotes unquantifiable fitness effects. Letters show wildtype bases, and the 3 mutations at each position are ordered alphabetically, as in **B**. Single promoter mutants make up the right-most column (*araA*) and top row (*araB*). Inducer concentrations were: 20 ng/ml aTc and 30 μM IPTG (Env_1_); 5 ng/ml aTc and no IPTG (Env_2_); 200 ng/ml aTc and no IPTG (Env_3_). **(B)** Fitness changes when an allele of one promoter is added to alleles of the second promoter. Large points indicate the “background” promoter is wildtype. Red, blue and grey points indicate positive, negative and non-significant fitness changes, respectively. Red, blue and grey rectangles indicate, in that environment, an allele can be beneficial but never deleterious, deleterious but never beneficial, or both beneficial and deleterious. G7A of P_LtetO-1_-*araA* (*) is the only allele conferring a qualitatively consistent fitness effect (beneficial) across all backgrounds and environments.

To further characterize how the effects of mutations in one gene depended on the allele present at the other gene, we computed epistasis (*17*) for all mutation pairs in each environment. Epistasis evaluates quantitatively and qualitatively how the log fitness of a double mutant deviates from the sum of that of the constituent single mutants (Fig. 3A, Fig. S6A). Epistasis was found to be pervasive (89%, 39% and 81% of pairs in Env_1-3_, respectively), heterogeneous and environment-dependent. A trend of antagonism reported for several other systems (*18*) was recovered between pairs of individually beneficial (negative epistasis in 89%, 72% and 100% in Env_1-3_, respectively) and individually deleterious (positive epistasis 100% (1/1), 97% and 98%, respectively) mutations, while interactions between a beneficial and a deleterious mutation could be mostly positive or mostly negative, depending on the environment and on which gene carried the beneficial/deleterious mutation. This epistatic diversity extended to individual mutation pairs, with more than 20% interacting both positively and negatively across environments (Figs. S6B-C). Notably, sign epistasis, an extreme interaction which occurs when the sign of a mutation effect changes in the presence of a second mutation (Fig. 1D), represented 31% of significant interactions in Env_1_, 17% in Env_2_ and 34% in Env_3_.

**Fig. 3.**
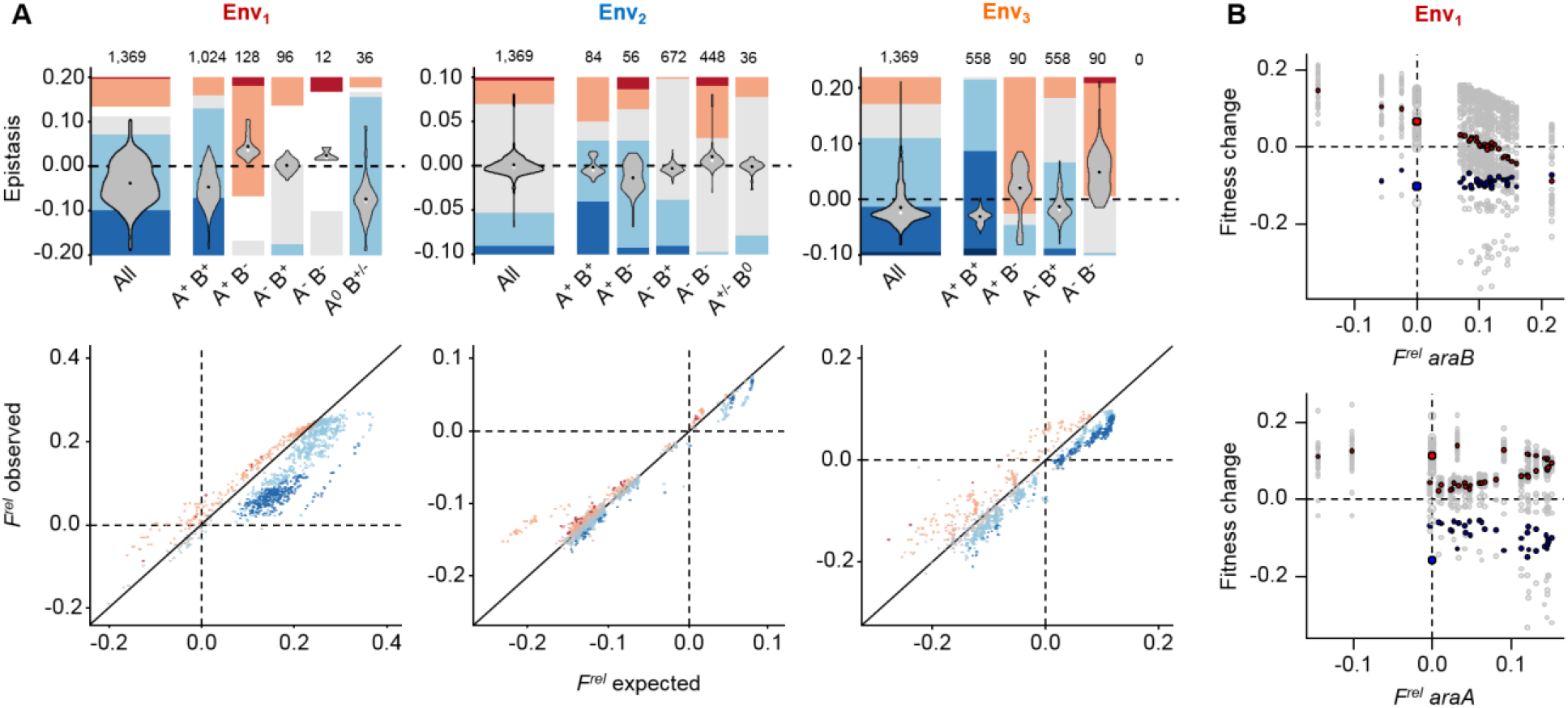
Strength, types and trends of epistasis across environments. **(A)** Violins show epistasis for different kinds of mutation pairs (white point - median; black point - mean). Mutation pairs may contain mutations that are individually both beneficial (A^+^ B^+^), both deleterious (AB^−^) or mixed (A^+^ B^−^ and A^−^ B^+^), or one of which confers an undetectable effect (A^0^ B^+/−^ and A^+/−^ B^0^). The number of each such pair is provided. Stacked bars show fractions of different epistasis types (colors as Fig. 1D, with white where epistasis could not be computed). Scatterplots show fitness of double mutants against that expected if mutation effects combined additively. Points colored as in Fig. 1D. **(B)** Relationship between background fitness and the fitness change induced by mutations in the second promoter, in Env_1_. Top: *araA* promoter mutations added to existing *araB* promoter mutations; bottom: inverse case. Colored points highlight particular alleles. Top: P_LtetO-1_-*araA* alleles T2C (red) and G7C (blue). Bottom: P_LtacO-1_-*araB* alleles T1A (red) and C11A (blue). Large points show effects in the wildtype background.

Confronted with such a variety of interactions, we asked whether they might be understood simply in terms of the quantitative fitness effects of the interacting mutations, as has been found for some other mutation sets (*19*). We found that the effects of individual mutations were weakly predictive of the type and value of epistasis they exhibited with mutations at the second promoter (Fig. 3A scatterplots). In all environments, there was a significantly negative correlation between the sum of individual fitness effects and the value of epistasis (Pearson’s r = −0.36, −0.37, −0.51 in Env_1-3_, respectively; p < 2.2×10^−16^ for all), a trend of diminishing returns that appears common across experimental systems (*19–22*) (Fig. S7A). However, when the two genes were considered separately, the relationship between individual fitness effects and epistasis was found to be markedly different between *araA* and *araB:* the negative correlation was stronger for P_LtetO-1_-*araA* mutations being added to existing P_LlacO-1_-*araB* mutations than for the inverse case (Figs. S7B-C; Pearson’s r = −0.67, − 0.73, −0.63 in Env_1-3_, p < 2.2×10^−16^ for all, vs. 0.12, −0.20 and −0.34, p < 1.6×10^−5^ for all), in which the correlation can even be positive, an extremely rare trend in existing studies (*19*). Moreover, we found that the average trend was in some cases strikingly non-monotonic (Figs. S7B-C), revealing that different alleles of a particular promoter can cause similar fitness changes on their own but interact very differently with alleles at the second promoter.

The relationship between individual mutation effects and epistasis was further complicated by the fact that it could be different for different alleles of the same promoter. For example, in Env_1_, the numerous beneficial P_Lteto-1_-*araA* mutations caused the average negative trend with P_LlacO-1_-*araB* background fitness, while the rare deleterious P_Lteto-1_-*araA* mutations showed no such trend (Fig. 3B, top panel). For individual P_LlacO-1_-*araB* mutations in P_Lteto-1_-*αrαA* backgrounds, the relationship was consistently non-monotonic, but had a different average direction for individually beneficial or deleterious alleles (Fig. 3B, bottom panel). Moreover, the trend for a given allele could vary greatly with the environment (Figs. S7B-C). These results demonstrate that genes interacting simply through their common participation in a linear pathway can exhibit complex, allele- and environment-dependent trends of epistasis. The smooth patterns exemplified by Fig. 3B, however, suggest that they may in principle be understood from an underlying phenotypic mechanism.

To this end, we constructed a quantitative model of the metabolic pathway, where fitness results from a balance between the benefit of flux (*23*) and the costs of intermediate toxicity (*14, 24, 25*) and AraA and AraB protein expression (*26–28*). Log fitness was computed as 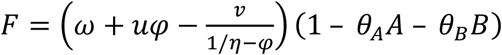, with *ω* a basal growth rate, *u* and *v* terms describing the catabolic benefit and toxicity cost of pathway flux (*φ*), *A* and *B* the cellular activity of the two enzymes, and θ the cost of enzyme expression. Flux depended on AraA and AraB activities as 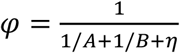 (*25, 29*).

Each promoter mutation was then characterized as a change in the activity (*via* expression) of AraA or AraB. Because most mutations lay outside of the repressor binding sites governing promoter inducibility (Fig. 1B), the fold-change in activity caused by each mutation was kept constant across inducer environments. Parameters describing the fitness function, wildtype activities in the 3 environments and expression effects of individual mutations were then optimized to fit the observed data (Data S2; Fig. S8A).

The fitted model is in excellent agreement with our data, yielding r^2^ values of 0.98 between experimental and simulated fitness effects and 0.82 between experimental and simulated epistasis coefficients (Fig. 4A-B; Fig. S8B-C; see Fig. S9 for more minimal models).

**Fig. 4.**
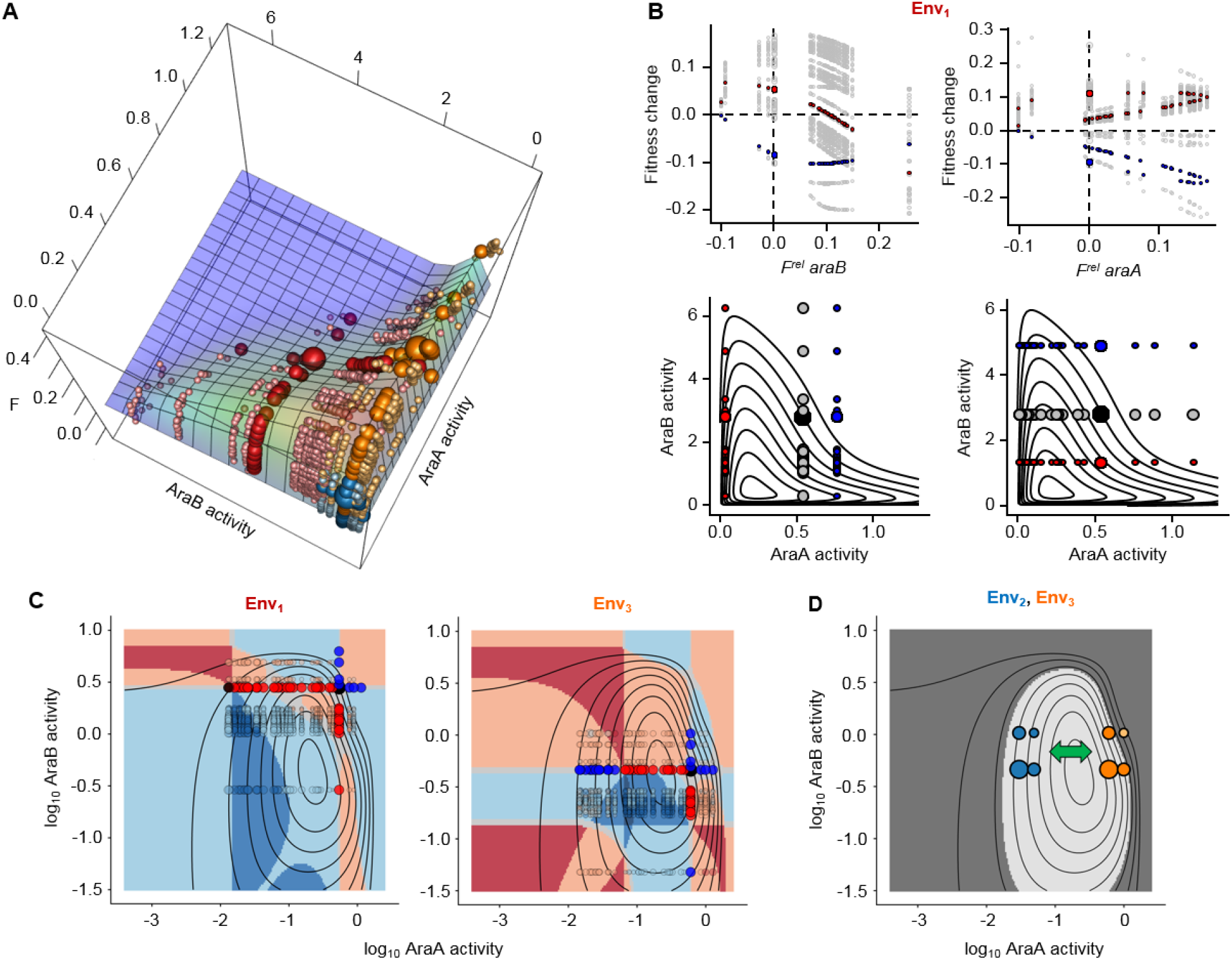
Mechanistic basis of heterogeneous, environmentally dependent epistasis. **(A)** Fitted activity-fitness model. Spheres are positioned according to predicted activity levels and observed *F^rel^* (Env_1-3_ – red, blue, orange). Three largest spheres are wildtype, intermediatesized spheres are single mutants, small pale spheres are double mutants. **(B)** Upper plots recapitulate Fig. 3B. Lower plots show highlighted genotypes within fitness landscape (black point is wildtype; other large points are single mutants, grey for the gene considered as carrying the “background” alleles). **(C)** Fitness surface on log activity scale, colored by predicted intergenic epistasis type (colors as Fig. 1D; determined as non-significant (grey) if magnitude < 0.005). Large black point is wildtype. Smaller, opaque blue, red and black points are single mutants, colored by observed *F^rel^* (deleterious, beneficial and neutral, respectively). Transparent points are double mutants, colored by observed epistasis type and sized by epistasis strength. **(D)** Dark grey marks area below a hypothetical disease threshold (40% of maximum fitness). Points are four genotypes in Env_2_ (blue) and Env_3_ (orange): wildtype (largest), C11A of P_LtetO-1_-*araA* and G7T of P_LtacO-1_-*araB* (intermediate size), and the resulting double-mutant (smallest). Green arrow represents a change in activity levels caused by non-genetic factors like ageing or environment. A disease state results here from one combination of alleles and environment (pale orange).

Notably, the model is capable of recapitulating the diverse and complex trends of epistasis seen in the data. In particular, we find that the non-monotonic relationships between singlemutant fitness and the fitness impact of alleles at the second promoter are well explained by the single mutants lying at two sides of a phenotypic optimum (Fig. 4B). Such overshooting, which is also the cause of sign epistasis (Fig. 4C) (*30*), is relatively common in our dataset, mostly because L-ribulose-5-phosphate toxicity results in an optimum in the flux-fitness relationship (*24, 25*) (Fig. S10). Two alleles of the same gene may thus result in similar fitness changes individually but cause substantially different expression levels and fluxes, resulting in different responses to mutations at the second gene. This is principally due to enzymes possessing different degrees of flux control on each side of the optimum, with lower levels of one resulting in the second having less control.

The model reveals how the biology underlying a linear pathway can result in heterogeneous, environmentally dependent intergenic interactions. When fitness depends solely on flux (*23, 25*), the nature of epistasis should be guaranteed by pathway topology alone (*25*). Under the slightly more complex selection pressure resulting from metabolite toxicity (*24, 25*) and gene expression costs, however, interactions can be both positive and negative. We find that epistatic categories form several localized zones over the fitness landscape, their size and position dependent on the wildtype phenotype, controlled here by the environment (Fig. 4C; Fig. S11). Encouragingly, these zones are generally large and orderly enough to be predictable, but only through knowledge of the underlying landscape and the position of the relevant genotypes within it.

The importance of this knowledge becomes immediately apparent when considering the existence of a disease threshold (Fig. 4D). The two alleles shown can lead to disease, but only when they co-occur, and only in one particular environment. The model thus provides a mechanism by which potential physiological defects can be manifested, aggravated or alleviated by particular combinations of alleles and environments (*1–7, 9*). Insight into intergenic fitness landscapes for other biological systems, and for genes connected by more complex topologies, will be indispensable for progress across medicine, bio-engineering and evolution.

## Acknowledgments

We thank A. Birgy, A. Decrulle, I. Matic, M. Deyell, A. Soler and D. Mazel for providing genetic material and technical advice, A. Baron, J. Chatel, A. Bridier-Nahmias and the CRI cytometry facility for technical assistance, and L.-M. Chevin, B. Gaut, E. Denamur and C. Landry for critical reading of the manuscript. MiSeq sequencing was performed using equipment provided by the Genetics Department of Bichat-Claude Bernard Hospital.

## Author contributions

H.E.K., P.N. and O.T. conceived the idea for the experiment; H.E.K. and O.T. designed the experiments; H.E.K., C.E., A.C., A.E.C., M.A.M., G.G. and H.L.N. performed the experiments; H.E.K., P.N. and O.T. performed the analyses; H.E.K., P.N. and O.T. wrote the paper.

## Data and materials availability

All genotype fitness estimates, along with their bootstrap 95% CIs and the number of replicates used to compute them, are provided in Supplementary Table 6. Raw and processed sequencing data has been submitted to GEO (accession number GSE115725). Custom code used in this study is available from the authors upon request.

## Supplementary Materials

Materials and Methods

Figures S1-S11

Tables S1-S5

References (*31*-*61*)

